# rVSV-EBOV vaccination protects ferrets from lethal Bundibugyo virus disease

**DOI:** 10.64898/2026.08.24.746878

**Authors:** Jordan Wight, Guodong Liu, Michael Chan, Sarah J. Medina, David Lu, Wenguang Cao, Samantha J. Krosta, Kevin Tierney, Kimberly Azaransky, Logan Banadyga

## Abstract

An uncontrolled and rapidly growing outbreak of Bundibugyo virus (BDBV) is currently gripping the Democratic Republic of the Congo and threatening health security across Central Africa. There are no available BDBV-specific vaccines, although emerging evidence suggests that the Ebola virus-specific vaccine, rVSV-EBOV (also known by its tradename ERVEBO), may offer cross-protective immunity. To directly address this question, we evaluated the efficacy of rVSV-EBOV in the uniformly lethal ferret model of BDBV infection. All vaccinated animals survived BDBV challenge and exhibited minimal clinical signs of infection, presumably as a result of a moderate—but protective—humoral immune response. These findings provide critical evidence further supporting the cross-protective efficacy of rVSV-EBOV, and they suggest a potential role for this vaccine in mitigating the ongoing BDBV outbreak.

## INTRODUCTION

On 15 May 2026, the Democratic Republic of the Congo (DRC) declared an outbreak of Bundibugyo virus (BDBV)—a filovirus closely related to Ebola virus (EBOV)—following laboratory confirmation of 8 cases, along with the identification of 246 suspected cases and 80 suspected deaths in the country’s Ituri Province (*1*). Two days later, the World Health Organization (WHO) declared this outbreak a Public Health Emergency of International Concern (PHEIC), reflecting, in part, fears over the spread of disease to Uganda and the significant uncertainty surrounding the true scope of the outbreak (*2*). Subsequent investigations suggest that the outbreak had begun months earlier, likely as a result of a new zoonotic spillover event (*3*, *4*), with hundreds of additional suspected cases identified between mid-January and mid-May (*5*). As of 12 August 2026, the outbreak has expanded to six provinces within the DRC, and cases of Bundibugyo virus disease (BVD) have been reported in 54 health zones, with 4,665 confirmed infections and 2,184 deaths (*6*). Twenty cases and two deaths have been reported in Uganda, and one case was exported to France (*6*). Although this represents only the third known BDBV outbreak, it is the largest BDBV outbreak to date, the largest filovirus outbreak in DRC, and the second largest filovirus outbreak on record.

There are currently no clinically approved therapeutics or medical countermeasures against BDBV. While multiple filovirus vaccines have been developed, with several having gone through clinical evaluation, only the recombinant vesicular stomatitis virus (rVSV)-based vaccine encoding the EBOV glycoprotein (rVSV-EBOV; tradename ERVEBO; Merck, https://www.merck.com) is currently licensed and clinically approved for use in humans. Multiple studies in animal models have shown that vaccines targeting one filovirus glycoprotein (GP) may provide some degree of cross-protection against other filoviruses, presumably due to the similarity of their GPs (*7–11*). Two studies in non-human primates (NHPs) vaccinated with rVSV-EBOV found partial cross-protection against lethal BDBV infection, with most animals surviving the infection (*8*, *11*). However, these findings were confounded by small group sizes, different vaccination schemes, and the fact that BDBV infection in NHP models is not uniformly lethal.

Ferrets have emerged as a promising filovirus animal model, particularly as they recapitulate all major hallmarks of filovirus disease and are highly susceptible to most wildtype filoviruses (*12*). Unlike the NHP models, BDBV infection in ferrets causes uniformly lethal disease (*13*, *14*) making these animals a valuable model for evaluating the efficacy of medical countermeasures, including vaccines (*15*). Recently, our group showed that ferrets vaccinated with rVSV-EBOV generate a robust IgG response that is cross-reactive with BDBV GP, an important pre-requisite for potential BDBV cross-protection (*16*). Given this evidence, the mixed results from earlier studies in NHPs, and the rapidly worsening epidemic in the DRC, we sought to evaluate the protective efficacy of the rVSV-EBOV vaccine against BDBV infection in the uniformly lethal ferret model.

## RESULTS

### rVSV-EBOV completely protected ferrets from lethal BDBV disease

One group of ferrets (*n*=6) received a single dose of rVSV-EBOV 28 days prior to BDBV challenge, while a second group (*n*=6) received a prime dose 28 days prior to challenge followed by a boost dose 14 days prior to challenge (**Table S1**). One control group (*n*=2) received saline instead of vaccine 28 days prior to challenge, while a second control group (*n*=2) did not receive saline and was challenged with BDBV 14 days after the first cohort (**Table S1**). All vaccinated animals were protected from a lethal dose of BDBV and survived until the end of the study at 29 days post-infection (DPI), whereas all control animals succumbed to the infection ( (1, *n* = 16) = 60.4, *p* = 8 x 10^-15^; **Fig. 1A**). Most vaccinated animals showed no overt signs of disease, other than a mild, transient fever that was observed in 5 of 12 ferrets (**Fig. 1B-D**). Although one animal from the prime-boost group (405M) lost nearly 10% body weight by 8 DPI and experienced a mild fever, the animal appeared otherwise healthy throughout the study. Conversely, unvaccinated control animals met the humane endpoint criteria and were euthanized on either 7 or 8 DPI (**Fig. 1A**). Ten historical control animals, which had all been inoculated with the same dose of BDBV, also succumbed to disease within the same time frame ( (1, *n* = 14) = 0.3, *p* = 0.6;**Fig. 1A**). Disease in the control animals was severe, with elevated body temperatures and weight loss accompanying severe clinical signs of disease, including evidence of hemorrhage in some animals (**Fig. 1B-D, Table S1**).

**Fig. 1.**
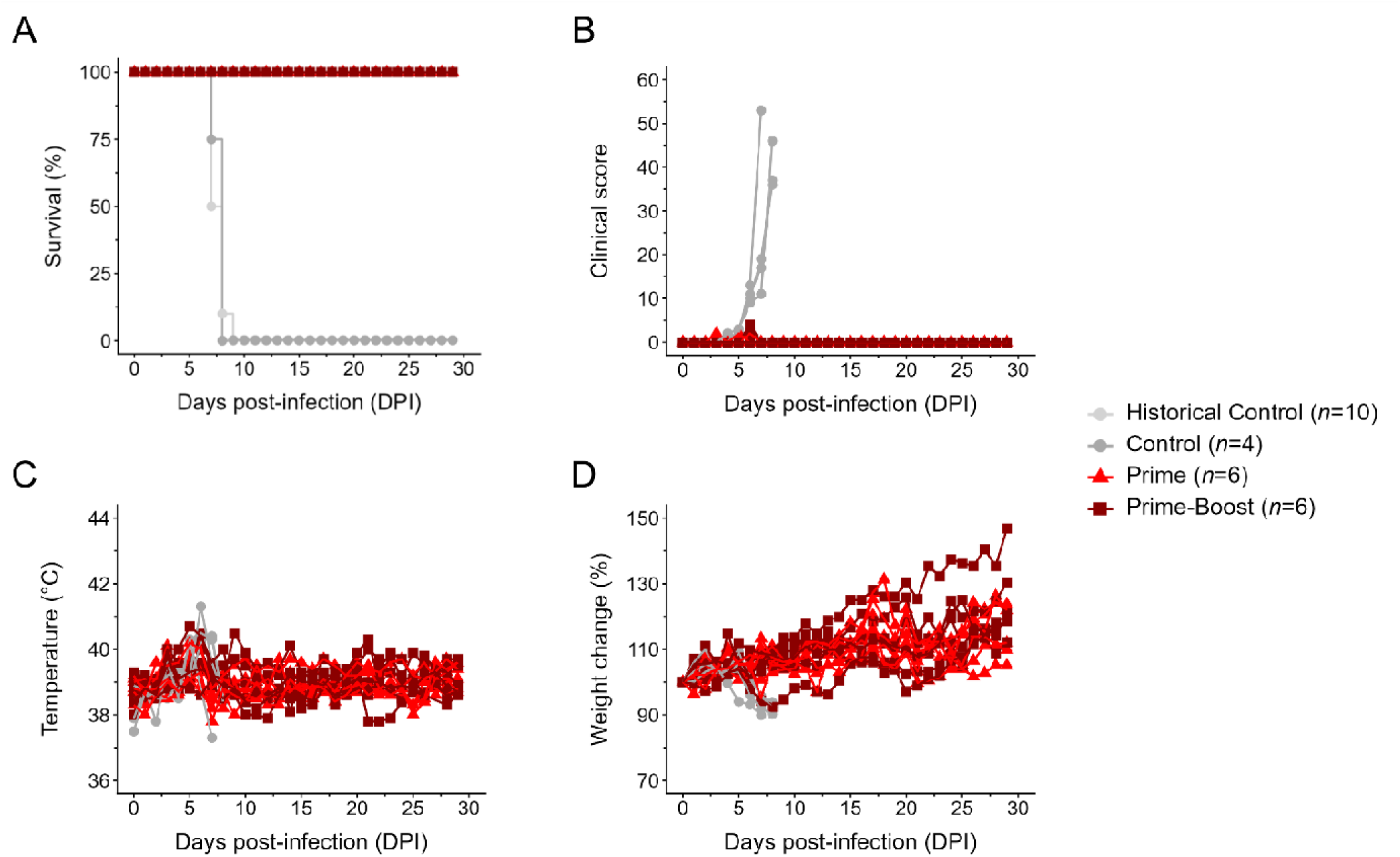
Ferrets vaccinated with rVSV-EBOV are protected from Bundibugyo virus disease. Following vaccination with rVSV-EBOV and inoculation with a lethal dose of BDBV, animals were monitored daily for survival (**A**), clinical score (**B**), body temperature (**C**), and body weight (**D**). Survival data for ten historical control animals which had all been inoculated with the same dose of BDBV are included in (**A**) in light gray.

Control animals also experienced changes in blood biochemistry typical of filovirus disease, with large increases in levels of alanine aminotransferase, alkaline phosphatase, total bilirubin, and blood urea nitrogen indicating significant liver and kidney damage (**Fig. 2**). Although two animals from the prime-boost group (405M and 499M) exhibited transiently elevated levels of alanine aminotransferase (at 6 and 9 DPI), levels of the other three analytes remained at or near baseline levels for all other vaccinated animals (**Fig. 2**). Increases in amylase, creatinine, phosphorus, and potassium levels were also observed in the control animals but not the vaccinated animals (**Fig. S1**). Neither control nor vaccinated animals showed dramatic changes in blood cell counts, although there was an apparent downward trend in platelet count observed for the control animals (**Fig. S2**).

**Fig. 2.**
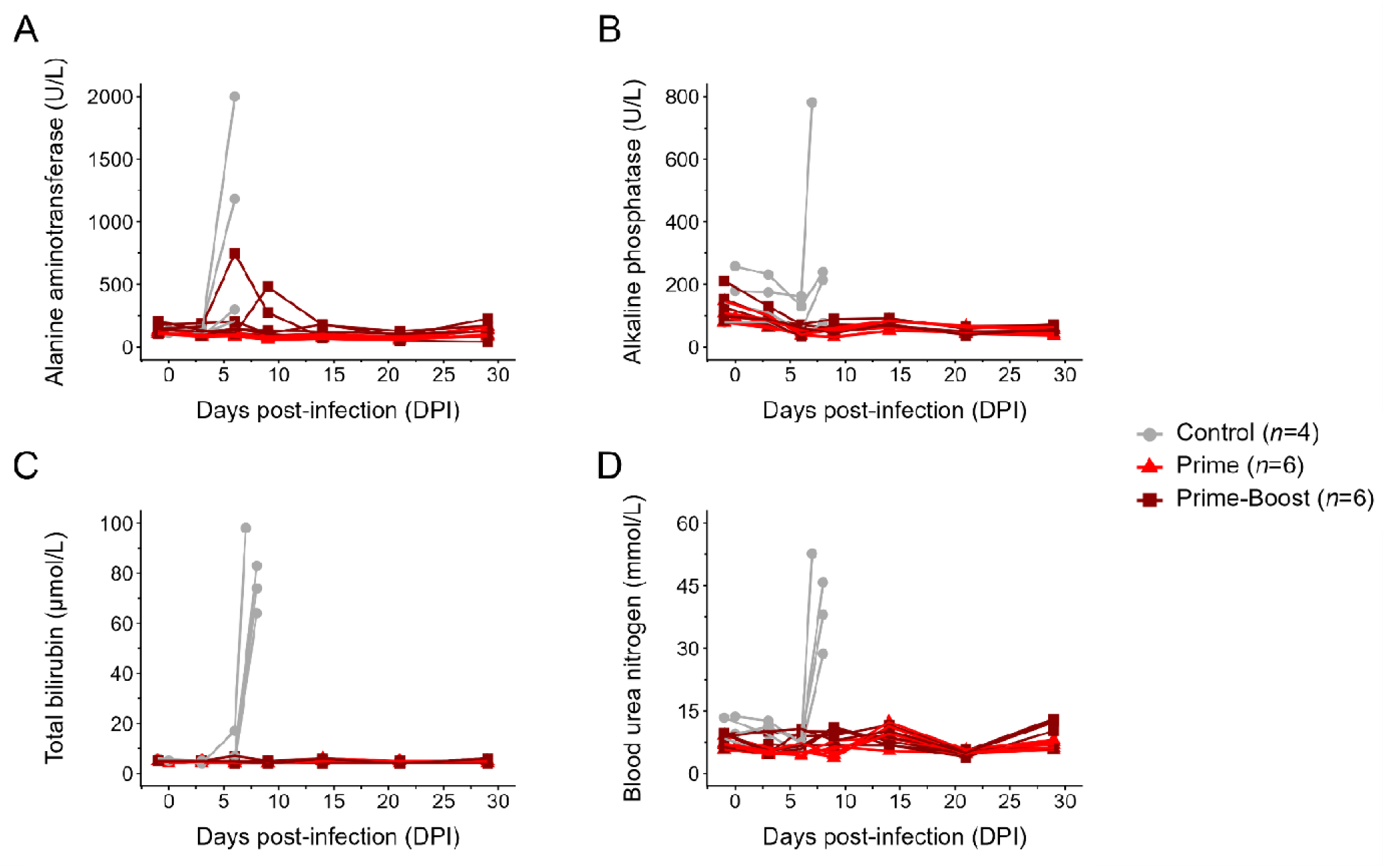
Vaccinated ferrets exhibit normal liver and kidney function. Alanine aminotransferase (**A**), alkaline phosphatase (**B**), total bilirubin (**C**), and blood urea nitrogen (**D**) levels were assessed in the blood of all ferrets after BDBV inoculation.

### Vaccination did not elicit sterilizing immunity

Despite the absence of overt clinical disease in most vaccinated animals, low to moderate levels of viral RNA were detected in the blood of all animals (**Fig. 3A**). Viral RNA levels peaked in the vaccinated animals on 6 DPI, with amounts less than 146 copies/µl (equivalent to a Ct value of 28), and declined to undetectable levels in most animals by 14 DPI. In the control animals, viral RNA levels were dramatically higher at 6 DPI and peaked at the terminal time points, around ∼630,000 copies/µl of blood (equivalent to a Ct value of 15). No infectious virus was detected in the blood of the vaccinated animals at 6 DPI, whereas relatively high levels of infectious virus were detected in the blood in 3 of the 4 control animals at the same time point (**Fig. 3B**). While viral RNA was detected in the nasal and oral swabs of all unvaccinated animals, low levels were detected in only a subset of vaccinated animals, with values once again peaking at 6 DPI (**Fig. 3C, E**). Low levels of infectious virus were detected in the nasal swabs of 6 vaccinated animals and in the oral swabs of 4 animals at 6 DPI, while 3 of 4 control animals exhibited similarly low levels of infectious virus (**Fig. 3D, F**). All but one vaccinated animal had detectable viral RNA in their rectal swabs, with amounts that decreased rapidly and were no longer detectable by the end of the study (**Fig. 3G**). Infectious virus was not detected in any of these samples at 6 DPI (**Fig. 3H**). These findings demonstrate that rVSV-EBOV vaccination did not confer sterilizing immunity against BDBV and that at least some of the vaccinated animals were shedding low levels of infectious virus.

**Fig. 3.**
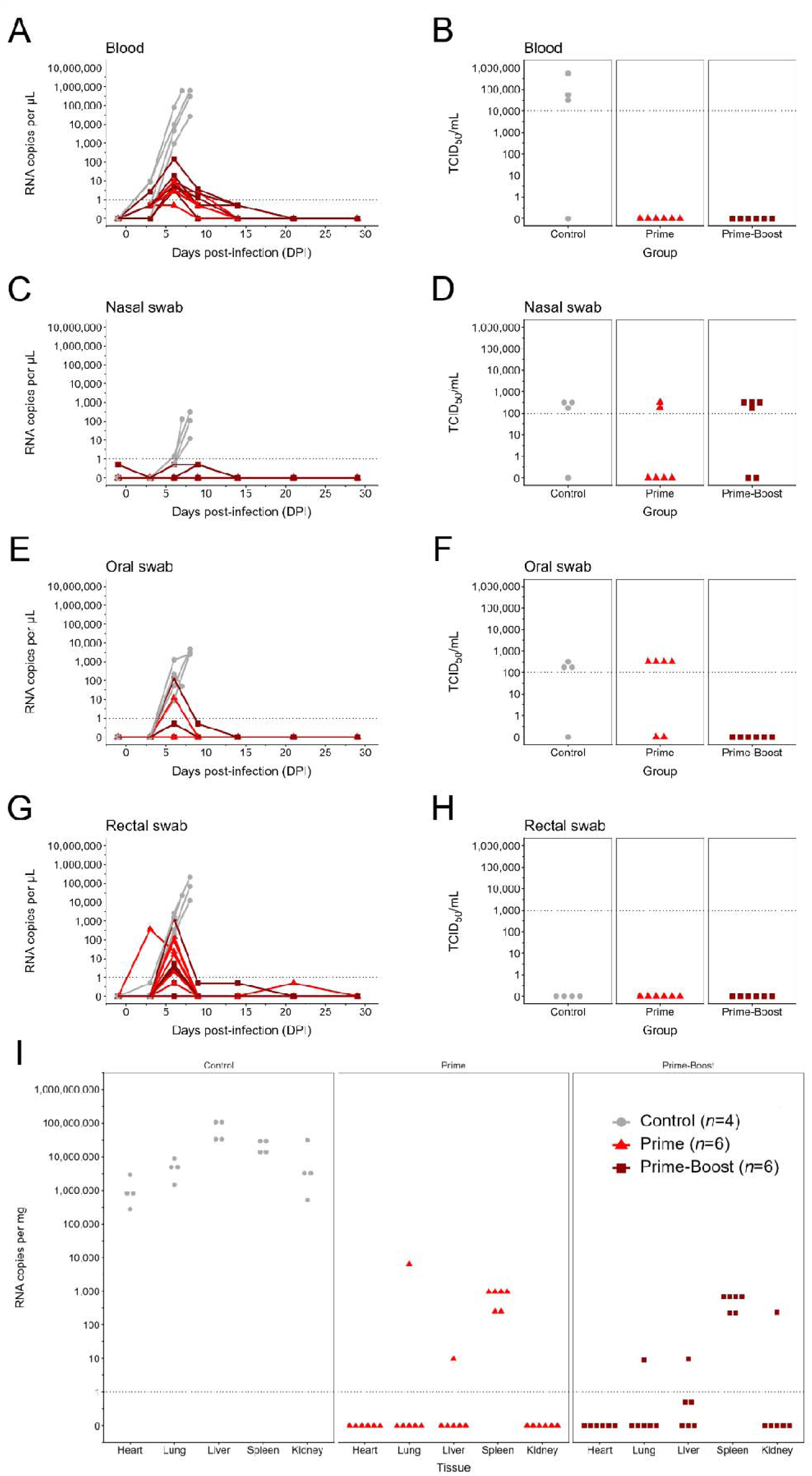
Viral RNA and infectious virus titers in blood, swab, and tissue samples. Viral RNA levels were quantified by RT-qPCR in blood (**A**), nasal swab (**C**), oral swab (**E**), rectal swab (**G**), and tissue samples (**I**). Levels of infectious virus were quantified via TCID_50_ assay in blood (**B**), nasal swab (**D**), oral swab (**F**), and rectal swab (**H**) samples collected at 6 DPI. The lower limit of detection of the RT-qPCR assay are represented as a dashed line at 1 RNA copy per µL or per mg. The lower limit of detection of the TCID_50_ assays are represented as a dashed line at 10,000 for blood (**B**), 100 for nasal and oral swabs (**D**, **F**), and 1,000 for rectal swabs (**H**).

At the terminal timepoints, all control animals showed high levels of viral RNA in the heart, lung, liver, spleen, and kidney (**Fig. 3I**). No viral RNA was detected in the majority of tissues from the vaccinated animals, with the exception of the spleen, in which low levels of viral RNA (less than ∼1,000 copies/mg of tissue; equivalent to a Ct value of 29) were detected in all animals (**Fig. 3I**).

### Vaccination induced cross-protective antibodies against BDBV

All vaccinated animals mounted a robust IgG response to EBOV GP with a lower level of reactivity to BDBV GP, as measured by indirect ELISA. For sera collected 27 days following vaccination (i.e., 1 day before BDBV challenge or -1 DPI), the geometric mean reciprocal endpoint titres for EBOV GP were 25,600 (range 6,400 to 51,200) in the single dose group and 36,204 (range 25,600 to 51,200) in the prime-boost group (**Fig. 4A**). Among the same animals, the rVSV-EBOV vaccine also induced antibodies that cross-reacted with the BDBV GP, although these levels were considerably lower, with titres of 898 (range 200 to 3,200) for the single dose group and 2,263 (range 1,600 to 6,400) for the prime-boost group. There was no statistically significant difference in anti-EBOV GP titres (*p* = 0.663) or anti-BDBV GP titres (*p* = 0.176) between animals that received one or two doses of rVSV-EBOV (**Fig. 4A**). By 29 DPI, BDBV-specific IgG titres increased in all vaccinated animals, indicating that the animals mounted an immune response following virus challenge. Antibody reactivity against BDBV GP increased significantly to a mean titre of 18,102 (range 6,400 to 25,600) in the single dose group and to 20,319 (range 6,400 to 51,200) in the prime-boost group (*p* = 0.004 and *p* = 0.006, respectively). Increases in reactivity against EBOV GP were also observed following challenge, with a modest, but not statistically significant, increase (*p = 0.214)* to 45,614 (range 25,600 to 51,200) in the single dose group and a significant increase (*p* = 0.02) to 91,228 (range 51,200 to 204,800) in the prime-boost group. Notably, the mean titre of the prime-boost group was significantly higher (*p* = 0.02) than the titre of the single dose group at 29 DPI, likely due to rising IgG levels following the second vaccine dose. Sera from unvaccinated control animals had no reactivity against EBOV or BDBV GPs at either -1 DPI or their terminal timepoints.

**Fig. 4.**
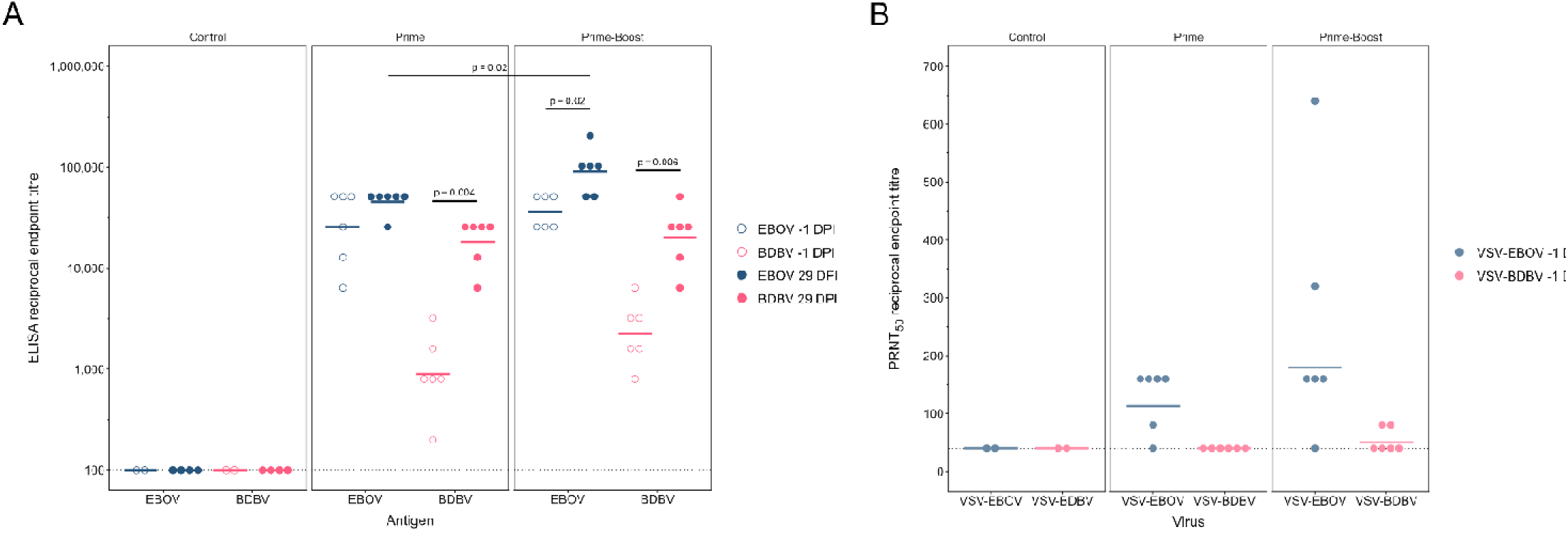
Vaccinated ferrets mount a cross-reactive humoral immune response to Bundibugyo virus. Gamma-irradiated serum was assessed for antibodies reactive against EBOV GP (blue) and BDBV GP (pink) for samples collected at -1 DPI (empty circles) and 29 DPI (vaccinated animals) or terminal time points (controls animals) (filled circles) (**A**). Serum collected at -1 DPI was also assessed for neutralization activity against rVSV-EBOV (light blue) and rVSV-BDBV (light pink) (**B**). Horizontal coloured lines represent the geometric mean reciprocal endpoint titers. The dashed line at a reciprocal endpoint titer of 100 (**A**) and 40 (**B**) represent the lower limit of detection of each assay.

Plaque reduction neutralization tests (PRNTs) using pseudotyped rVSVs were also conducted on sera collected 27 days following vaccination. Overall, the PRNT_50_ titres were much lower than the IgG titres measured by ELISA, and they were higher against rVSV-EBOV than rVSV-BDBV (**Fig. 4B**). Against rVSV-EBOV, the geometric mean reciprocal PRNT_50_ titre was 113 (range <40 to 160) in the single dose group and 180 (range <40 to 640) in the prime-boost group. Against rVSV-BDBV, no sera from animals that received a single dose had a measurable neutralization titre against rVSV-BDBV (i.e., <40), and only sera from two animals in the prime-boost group had any neutralizing activity at titres of 80 (geometric mean 50, range <40 to 80). Again, sera from unvaccinated controls had no detectable neutralization against either rVSV-EBOV or rVSV-BDBV.

## DISCUSSION

The ongoing BDBV outbreak in the DRC is already unprecedented in size and scope. Confronting a rapidly growing epidemic caused by a previously “neglected” filovirus (*17*) within the context of an unstable local and international geopolitical landscape, requires consideration of all public health tools available. In the absence of BDBV-specific medical countermeasures, rVSV-EBOV (i.e., ERVEBO) has been proposed as a potentially effective—albeit interim— solution that could quickly and relatively easily be deployed to mitigate the outbreak and begin saving lives (*18*). This position is supported by modelling efforts that suggest even a partially cross-protective vaccine could greatly reduce morbidity and mortality (*19*).

Several studies have generated evidence that points towards the potential utility of rVSV-EBOV against BDBV. Retrospective serological studies of humans vaccinated with rVSV-EBOV have demonstrated serological cross-reactivity against BDBV (*20–22*), providing support for the idea that rVSV-EBOV may confer some level of protection against BDBV. Anecdotal evidence from the current outbreak has also suggested that patients who had previously been vaccinated with rVSV-EBOV may fare better in the face of BVD than their unvaccinated counterparts (*18*, *23*). In conjunction with limited evidence suggesting that BDBV cross-protection may be achievable in NHPs (*8*, *11*), along with our observation that rVSV-EBOV elicits a cross-reactive IgG response in vaccinated ferrets (*16*), we sought to evaluate the cross-protective efficacy of rVSV-EBOV using the uniformly lethal ferret model of BVD.

Ferrets vaccinated with rVSV-EBOV and challenged with BDBV four weeks later were fully protected from lethal disease, even after receiving only a single dose of the vaccine. No pronounced clinical signs of disease were observed, and the overall clinical picture for all vaccinated animals for the duration of the study was largely benign. In contrast, the control animals experienced severe disease, with markedly high levels of viral RNA in blood, swab, and tissue samples, along with dramatic perturbations in hematology and blood biochemistry parameters, and obvious clinical signs of filovirus disease, including hemorrhagic manifestations. Despite protection from disease, vaccination did not provide sterilizing immunity, with virus shedding observed alongside evidence suggestive of viremia in many vaccinated animals. Viral RNA was detected in the blood of all vaccinated animals, peaking on 6 DPI, although infectious virus was not detected at the same time point. Whether infectious virus was present at levels below the limit of detection for this assay (i.e., 10,000 TCID_50_/mL) is unclear but cannot be ruled out. Viral RNA levels in the oral and nasal swabs, accompanied by infectious virus in some of the samples, indicates that at least a few of the animals were shedding infectious virus. While we did not detect infectious virus in the rectal swabs, even with consistent detection of low to moderate levels of viral RNA, we also cannot rule out the presence of virus below this assay’s limit of detection (i.e., 1,000 TCID_50_/mL). The absence of sterilizing immunity against BDBV in ferrets vaccinated with rVSV-EBOV reflects similar observations in NHPs (*8*) and suggests that heterologous vaccination in this context may prevent disease but not transmission.

The protection from BVD conferred by rVSV-EBOV was most likely mediated by the humoral immune response mounted by all vaccinated animals, in line with previous observations that non-neutralizing antibodies are critical to rVSV-EBOV-mediated protection against EBOV in animal models, including NHPs and humans (*24–28*). In addition to exhibiting high levels of IgG reactive against EBOV GP, vaccinated animals also exhibited lower, but presumably still functionally significant, levels of IgG reactive against BDBV GP. Importantly, we also observed a significant increase in BDBV GP-specific IgG levels in the surviving animals, indicating that they mounted their own humoral immune response to BDBV following challenge. Collectively, these findings suggest that the surviving animals would have robust protection against subsequent exposure to either EBOV or BDBV, which may have important implications in the large scale deployment of rVSV-EBOV during the current outbreak.

In contrast to the overall levels of IgG reactive against EBOV or BDBV GP, neutralizing antibody levels were substantially lower and notably absent in most animals against BDBV GP. Most animals exhibited low levels of neutralizing activity against EBOV GP, while neutralizing activity against BDBV GP was detected in only two animals from the prime-boost cohort. Nevertheless, the fact that all vaccinated animals survived lethal BDBV infection suggests that the neutralizing response was either unnecessary for protection in this context or undetectable using our assay. It is also worth noting that, despite the lack of a robust cross-neutralizing response, we saw no evidence of antibody-dependent enhancement (ADE) of infection. Although ADE of filovirus infection has been demonstrated *in vitro* (*29*), raising concerns that non-neutralizing antibodies could influence the outcome of heterologous vaccination (*8*, *30*), there is currently no compelling *in vivo* evidence to support this phenomenon. Unfortunately, our study did not examine the cell-mediated immune response to vaccination, so its contributions, if any, to heterologous protection remains unclear.

As the results of this study may be used to inform the public health response to the ongoing outbreak of BDBV in the DRC (*31*), at least two important considerations are warranted. First, we note that although ferrets are valuable and robust models of filovirus disease that accurately recapitulate many of the hallmark clinical signs observed in humans and NHPs, there may be significant—and unappreciated—differences between the cross-protective efficacy of this vaccine in different experimental model systems and humans. Whether and how immunity against BDBV differs in NHPs, humans, and ferrets is effectively unexplored. Second, the correlates of protection against BDBV remain poorly understood, in general, and almost all of what we do know has been largely extrapolated from studies with EBOV. This uncertainty therefore limits our ability to understand exactly what contributes to a protective immune response against BDBV, regardless of whether protection is elicited by a heterologous or homologous vaccine. Given these considerations, the data presented here cannot unilaterally validate the hypothesis that rVSV-EBOV will provide cross-protection against BDBV in humans, which is something that must ultimately be evaluated in Phase 3 clinical trials.

Effective medical countermeasures are urgently needed to address the rapidly worsening public health emergency caused by BDBV in the DRC. Although BDBV-specific vaccines would be the ideal solution, promising candidates remain in development and will require substantial pre-clinical and clinical evaluation. In the interim, rVSV-EBOV represents a safe, effective, clinically approved, and readily available vaccine for the related EBOV (*18*, *32*). Emerging evidence from animal and human studies indicates that rVSV-EBOV can elicit cross-reactive immunity against BDBV, and our findings presented here extend this evidence by demonstrating cross-protective efficacy of this vaccine in the ferret model. Whether this protection can be extended to humans remains unknown, but given the urgency of the outbreak, the potential for rVSV-EBOV to reduce morbidity and save lives warrants careful consideration.

## MATERIALS AND METHODS

### Animal ethics and biosafety statement

Animal work was conducted at the Canadian Science Centre for Human and Animal Health (CSCHAH), National Microbiology Laboratory of the Public Health Agency of Canada in Winnipeg, Manitoba. Animal use protocols were approved by the Institutional Animal Care Committee in accordance with guidelines from the Canadian Council on Animal Care. Work with rVSVs and filoviruses was conducted in biosafety containment level 2 and 4 laboratories, respectively, at the CSCHAH in accordance with standard operating protocols.

### Viruses and cells

A research-grade recombinant vesicular stomatitis virus (rVSV) expressing the EBOV variant Kikwit glycoprotein in place of the endogenous VSV glycoprotein (rVSV-EBOV) (*33*) was used as the vaccine, while BDBV (Bundibugyo virus/H.sapiens-tc/UGA/2007/Butalya-811250; GenBank accession # NC_014373) was used as the challenge virus. Vaccine and challenge virus stocks were mycoplasma negative, sequence confirmed, and stored in -80°C. Vero E6 (ATCC CRL-1586) cells were maintained in growth medium comprised of Dulbecco’s Modified Eagle’s Medium (DMEM; ThermoFisher Scientific, Cat # 11965092) supplemented with 10% heat-inactivated fetal bovine serum (FBS; Corning, Cat # 35-077-CV), 100 U/mL of penicillin and 100 µg/mL streptomycin (ThermoFisher Scientific, Cat # 15140122), and 2 mM L-glutamine (ThermoFisher Scientific, Cat # 25030081).

### Study design

Sixteen approximately 12-week old ferrets (*Mustela putorius furo*; 8 male and 8 female), implanted with an ID and temperature transponder, were purchased from Marshall BioResources. Animals were acclimatized for at least seven days before study commencement. All animals were monitored at least daily and were provided food and water *ad libitum*. All invasive manipulations, such as blood collection, swab sampling, vaccination, and infection, were performed under isoflurane anaesthesia.

Ferrets were randomly assigned to vaccination and control groups, with an equal number of males and females in each group. One group of six animals received a single dose (prime) of 1 x 10^6^ PFU of rVSV-EBOV 28 days before infection, while a second group of six animals received two doses (prime and boost) of vaccine at 28 and 14 days before infection. The vaccine was diluted in 0.9% saline to a total volume of 500 µL and delivered via intramuscular injection to the left (prime) or right (boost) quadricep. Control animals (*n*=2) received 500 µL of 0.9% saline 28 days before infection. A second group of control animals (*n*=2) was added to the study 14 days after the first cohort of animals were challenged with BDBV; these animals did not receive saline.

Twenty-eight days following the initial vaccination, animals were inoculated with a dose of 1,000 TCID_50_ of EBOV via the intramuscular (IM) route in a total volume of 500 µL divided equally between both quadriceps. The temperature and weight of each animal was measured daily for the 29 day period following challenge. On predetermined days (−1, 3, 6, 9, 14, 21, 29 days post-infection, DPI) or when an animal reached a clinical score requiring euthanasia, animals were anesthetized, given a physical examination by study personnel, and blood and nasal, oral, and rectal swabs were collected. At the end of the study, or when animals reached humane clinical endpoint, necropsies were performed and heart, lung, liver, spleen, and kidney tissue were collected for further analyses. Blood was collected from the jugular vein and transferred to either microtubes containing potassium EDTA (Sarstedt Inc, Cat # 41.1395.105), lithium heparin (Sarstedt Inc, Cat # 41.1393.105), or serum separation gel (Sarstedt Inc, Cat # 41.1378.005) depending on the intended application. For serum, serum separation tubes were spun at 3,500 g for five minutes, after which serum was removed and subsequently stored at - 80°C. Nasal, oral, and rectal swabs (Puritan, Cat 25-860 1PD) were collected, with the nasal swab pre-wetted in DMEM prior to sampling. Following collection, swabs were placed into separate vials containing 1 mL of plain DMEM.

### Blood Biochemistry and Hematology

Complete blood cell counts in EDTA-treated blood were performed using the VetScan® HM5 hematology analyzer (Zoetis Services LLC) to enumerate total white blood cells, lymphocytes, monocytes, neutrophils, and platelets. Blood biochemistry was analyzed using lithium heparin-treated blood on a VetScan® VS2 chemistry analyzer (Zoetis Services LLC) and VetScan® Comprehensive Diagnostic Profile reagent rotors. The following analytes were quantified: alanine aminotransferase, albumin, alkaline phosphatase, amylase, calcium, creatinine, globulin, glucose, phosphorus, potassium, sodium, total bilirubin, total protein, and blood urea nitrogen.

### RNA extraction and determination of viral RNA load

For liquid samples, 140 µL of EDTA-treated blood or swab supernatant was added to 560 µL of Buffer AVL (Qiagen, Cat # 19073) containing 10 µg/mL carrier RNA and incubated at room temperature for 10 minutes. 560 µL of 100% ethanol was then added for a total volume of 1,260 µL, and samples were removed from CL4 using approved protocols. For tissue samples, sections of heart, lung, liver, spleen, or kidney tissue were transferred to a vial containing 1,000 µL of RNAlater™ stabilization solution (Invitrogen, Cat # AM7021) and incubated at 4°C overnight. The following day, the vial was centrifuged, and RNAlater™ was removed, weighed, and 600µL of Buffer RLT (Qiagen, Cat # 79216) containing 0.01% -mercaptoethanol was added along with a single 5 mm stainless steel homogenization bead (Qiagen, Cat # 69989). Tissue was homogenized using a Bead Ruptor Elite Tissue Homogenizer (Omni International, Inc.) for 30 seconds at a frequency of 4 m/s. The amount of homogenate corresponding to 30 mg of tissue was transferred to a new tube, and the volume was brought up to 600 µL with RLT, mixed, and incubated at room temperature for 10 minutes. 600 µL of 70% ethanol was added for a total volume of 1,200 µL, and samples were removed from CL4 using approved protocols. RNA extractions were performed in CL2 using the MagMax™ Viral/Pathogen II Nucleic Acid Isolation Kit (Applied Biosystems, Cat # A48383) on a KingFisher Apex System (Thermo Scientific) using a modified protocol. Briefly, 970 µL of sample removed from CL4 was added to each well of the extraction block containing 20 µL of binding beads and 10 µL of proteinase K. A modified version of the standard extraction program was performed on the instrument, where a second wash with 500 µL of 80% ethanol was included. RNA was eluted in 60 µL or 50 µL of elution buffer for liquid and tissue samples, respectively, and subsequently stored at -80°C.

RT-qPCR was performed to determine viral RNA loads using TaqPath™ 1-Step Multiplex Master Mix (Applied Biosystems, Product# A28523) on a QuantStudio™ 5 Real-time PCR System (Applied Biosystems) and analyzed using the QuantStudio™ Design & Analysis Software version 1.5.3 (Applied Biosystems). The RT-qPCR targeting the BDBV L (polymerase) gene was prepared in a total reaction volume of 20 µL consisting of 5 µL of 4X master mix, 0.1 µL of 20 µM BDBV_L_F1 (5’-CCGAGAAAATCCACCAGAAG-3’), 0.1 µL of 20 µM BDBV_L_F2 (5’-TCGGGAAGATCCCCCGGAAG-3’), 0.2 µL of 20 µM BDBV_L_R (5’-TGTTGRAGTCCCTCAATYCC-3’), 0.4 µL of 10 µM BDBV_L_P (5’-[FAM]-CCAAGCTCT[ZEN]TACCGTGGTCATCTTGG-[IBFQ]-3’), 9.2 µL of nuclease-free water, and 5 µL of RNA. Cycling was performed in standard mode, with parameters as follows: 53°C for 10 minutes, 95°C for 2 minutes, followed by 45 cycles of 95°C for 3 seconds, and 60°C for 30 seconds at which time fluorescent signal was detected. Thresholds were determined automatically by the instrument’s software based on a standard curve made from serially diluted *in vitro* transcribed RNA included on each assay plate, manually confirmed, and applied to each sample to determine the cycle threshold (Ct) value and corresponding number of RNA copies.

*in vitro* transcribed RNA control material was generated by amplifying a portion of the L gene flanking the primer binding regions used for the RT-qPCR from viral stock RNA, cloned into the pGEM T-easy vector (Promega, Cat # A1360), and transformed into Stellar Chemically Competent Cells (Takara, Cat # 636763). Clones were screened by PCR and confirmed by Sanger sequencing, and a clone with the amplicon of interest cloned in the reverse direction was selected and plasmid DNA extracted using the Monarch® Spin Plasmid Miniprep Kit (New England Biolabs, Cat # T1110). 2 µg of plasmid was linearized by digestion with SacI-HF® (New England Biolabs, Cat # R3156) and purified using the Monarch® Spin PCR & DNA Cleanup Kit (New England Biolabs, Cat # T1130). *in vitro* transcription was performed using the HiScribe® T7 High Yield RNA Synthesis Kit (New England Biolabs, Cat # E2040) with 1 µg of linearized plasmid as input according to the manufacturer’s instructions for short transcripts. Plasmid DNA was removed by digestion with RNase-free DNase I (New England Biolabs, Cat # M0303) then RNA was purified using the RNeasy MinElute Cleanup Kit (Qiagen, Cat # 74204). RNA purity was assessed using a NanoDrop™ One spectrophotometer (Thermo Scientific) and quantified using the Qubit™ RNA High Sensitivity Assay Kit (Invitrogen, Cat # Q32852) on a Qubit™ 4 fluorometer (Invitrogen) and stored at -80°C. RNA copy number was determined using the ssRNA mass moles converter (*34*), taking into account the transcribed RNA sequence and the RNA concentration/mass.

On the day of the RT-qPCR assay, *in vitro* transcribed RNA was thawed, re-quantified by Qubit™, and used to calculate the RNA copy number of the first dilution, after which a serial ten-fold dilution was prepared in nuclease-free water in DNA LoBind® tubes (Eppendorf, Cat # 022431021). The prepared dilutions for the 11-point standard curve were included in duplicate on each assay plate along with no-template controls. To calculate the number of RNA copies present per µL of blood/swab supernatant or per mg of tissue, the fraction of the volume extracted out of the total volume removed from CL4 was used to calculate the amount of sample extracted. This was divided by the elution volume and multiplied by 5 based on the sample volume used for RT-qPCR. The quantity of RNA determined using the standard curve was divided by either 8.98 for liquid samples or 2.43 for tissue samples to calculate the number of RNA copies per µL of blood/swab supernatant or per mg of tissue, respectively. For the *in vitro* transcribed RNA standard curve, quantities of RNA were divided by 5 to calculate the number of RNA copies and are represented as either per µL or per mg, assuming a density of ∼1 g/cm^3^. Using data from all plates for each assay, the mean RNA copy number of the last dilution that provided amplification in all wells was determined as the limit of quantification (LOQ). For all samples that yielded a characteristic amplification curve but had an RNA amount less than the LOQ, these amounts were set as half of the LOQ and are displayed as such in all figures. For samples that yielded no amplification, the quantity is denoted as zero.

### Quantification of infectious virus

Blood samples collected on 6 DPI from surviving animals or the terminal timepoint for those that succumbed to the infection were assessed for the presence of infectious virus by median tissue culture infectious dose (TCID_50_) assays. VeroE6 cells were seeded in 96-well tissue culture plates. On the day of the assay, cells had reached a confluency of 90% and growth medium was removed and replaced with 100 µL of infection medium, prepared the same as growth medium but with 2% FBS. Serial ten-fold dilutions of EDTA-treated blood or swab supernatant were prepared in infection medium and 100 µL of each dilution was added to triplicate wells. Plates were incubated at 37°C with 5% CO_2_ and assessed on 14 DPI for cytopathic effects. The median TCID_50_ result was calculated for each sample using the Reed-Muench method (*35*).

### IgG ELISAs

EBOV GP-and BDBV GP-specific IgG levels were determined in gamma irradiated (5 Mrad) serum samples collected the day before challenge (−1 DPI) and at the terminal timepoint by indirect ELISA. Each serum sample was serial two-fold diluted (1:200 to 1:204,800) and tested in duplicate. Serum collected before vaccination was tested only at 1:200. ELISAs were performed largely as per (*16*), with minor modifications. Briefly, half-area high-binding 96-well assay plates (Corning, Cat # 3690) were coated using recombinant transmembrane domain-deleted glycoproteins, specifically EBOV (IBT Bioservices, Cat # 0501-015) or BDBV (IBT Bioservices, Cat # 0505-015); each serum sample was tested for antibodies against each of these recombinant glycoproteins. Recombinant proteins were diluted to 1 µg/mL in 50 mM carbonate-bicarbonate buffer pH 9.6. 30 µL of this solution was added to each well and plates were coated overnight at 4°C. The following day, the coating solution was removed and wells were blocked with 100 µL of 5% skim milk prepared in phosphate-buffered saline (PBS) pH 7.4 for 1 hour at 37°C. After removing the blocking solution, 30 µL of serum samples diluted as above in 2% skim milk in PBS were added to duplicate wells and incubated for 1 hour at 37°C. Plates were washed four times with 0.1% Tween-20 in PBS, and 30 µL of anti-ferret IgG (H+L) HRP secondary antibody (Novus Biologicals, Cat # NB7224) diluted 1:10,000 in 2% milk was added per well. After washing plates as above, 50 µL of 3,3’,5,5’-tetramethylbenzidine (TMB; Invitrogen, Cat # 002023) was added per well and incubated in the dark for 30 minutes. Color development was stopped by adding 50 µL of 0.16 M sulfuric acid to each well. The absorbance at 450 nm and 650 nm was measured using a Varioskan LUX Microplate Reader (Thermo Scientific) and the absorbance at 450 nm minus the absorbance at 650 nm was used to calculate the corrected absorbance. Positivity cut-off values were established as the mean plus three times the standard deviation of the 1:200 dilution of pre-vaccination serum collected from all ferrets. Endpoint dilution titres were determined as the highest dilution that exceeded the cut-off value.

### Plaque Reduction Neutralization Test (PRNT)

Serum samples collected the day prior to challenge (−1 DPI) were evaluated for the presence of neutralizing antibodies to rVSV-EBOV and rVSV-BDBV by PRNT assay. Serum was heat-inactivated at 56°C for 30 minutes, spun down at 8,000 g for five minutes, then serially diluted two-fold (starting at 1:40) in plain DMEM. Vaccine virus was diluted in plain DMEM to 667 PFU/mL, and 275 µL was added to the same volume of diluted serum to achieve final serum dilutions of 1:80 to 1:10,240. The serum and virus mixture was incubated for 1 hour at 37°C with 5% CO_2_, then 150 µL (containing 50 PFU of virus) was added to each well of a 24-well plate containing 90% confluent monolayers of Vero E6 cells. Plain DMEM was used as a negative control, and the above diluted virus mixed 1:1 with plain DMEM served as the positive control. After incubation for 1 hour at 37°C with 5% CO_2_, the inoculum was removed and overlaid with 0.5 mL per well of molten agarose overlay media cooled to 43°C comprised of 2% SeaPlaque® Agarose (Lonza, Cat # 50100) prepared in sterile water and mixed 1:1 with 2X Temin’s Modified Eagle Medium (MEM; ThermoFisher Scientific, Cat # 11935046) containing 4% FBS, 200 U/mL of penicillin and 200 µg/mL streptomycin, and 4 mM glutamine. At 3 DPI for rVSV-EBOV and 4 DPI for rVSV-BDBV, cell monolayers were fixed by adding 0.5 mL of 10% formalin containing crystal violet (0.2% *w*/*v*) and incubated at room temperature overnight. Fixative was removed, wells washed with water, and once dry, plaques were counted manually. The percent inhibition was calculated by dividing the mean number of plaques for each dilution by the mean number of plaques in the positive control. The highest dilution that resulted in a 50% inhibition was determined as the PRNT_50_. For samples where the PRNT_50_ titre was below the lower limit of detection (i.e., 1:80), a value of 1:40, being half the limit of detection, was assigned.

### Statistical analysis and data visualization

R v4.4.3 (*36*) was used to perform data manipulation, statistical analysis, and data visualization. A *p*-value < 0.05 was considered as significant for all tests. To evaluate differences in survival between vaccinated and control animals, log-ranked tests were performed using the survdiff function in the survival package v3.8-11 (*37*). To compare reciprocal endpoint titers between EBOV GP and BDBV GP antigens and pseudotyped virus neutralization titres between the prime and prime-boost regimens, Wilcoxon (W) rank-sum tests with continuity correction were performed. Additional R packages used included ggtext v0.1.2 (*38*), patchwork v1.3.2 (*39*), rstatix v0.7.3 (*40*), scales v1.4.0 (*41*), and those belonging to the tidyverse v2.0.0 (*42*) suite.

## Acknowledgements

We thank the Veterinary Technical Services team at the National Microbiology Laboratory (NML) for animal care in CL2 and technical assistance. We also thank Anders Leung and Yvon Deschambault for assistance with animal care in CL4.

## Author Contributions

Conceptualization: JW, LB

Data curation: JW, GL, MC, SM, DL

Formal analysis: JW

Investigation: JW, GL, MC, SM, DL, WC, SK, KT, KA, LB

Methodology: JW, GL, MC, LB

Software: JW

Visualization: JW

Resources: LB

Project administration: JW, LB

Supervision: LB

Funding acquisition: LB

Writing – original draft preparation: JW, LB

Writing – review and editing: JW, GL, MC, SM, DL, WC, SK, KT, KA, LB

All authors read and approved the final manuscript.

## Data availability

Data supporting the conclusions of this study can be found within the article or the supplementary information. Additional data are available from the corresponding author upon reasonable request.

### Supplementary Materials

Figs. S1 to S2 Table S1

## REFERENCES

1. CDC, Ebola Disease Outbreak in the Democratic Republic of the Congo and Uganda, Health Alert Network (HAN) (2026). https://www.cdc.gov/han/php/notices/han00530.html.

2. WHO, Epidemic of Ebola Disease caused by Bundibugyo virus in the Democratic Republic of the Congo and Uganda determined a public health emergency of international concern (2026). https://www.who.int/news/item/17-05-2026-epidemic-of-ebola-disease-in-the-democratic-republic-of-the-congo-and-uganda-determined-a-public-health-emergency-of-international-concern.

3. A. Amuri-Aziza, G. Luakanda-Ndelemo, A. Ayitewala, D. Jansen, P. Adroba-Tandele, A. Tebba, E. Kinganda-Lusamaki, S. Kanyerezi, P. Kaleebu, C. Ngandu, A. Muruta, D. Jjingo, S. Nabadda, R. Lumembe-Numbi, O. Ntumba-Tshitenge, M. Wayengera, I. Mugerwa, J. O. Otshudiema, J. Kyokushaba, D. J. Kyabayinze, P. Akil-Bandali, Ebola outbreak response consortium, R. Ola-Mpumbe, T. Bakutumba-Loweya, F. Cikaya-Kankolongo, D. Isengelo-Sikatenda, S. Kimbonza, J. Tete-Sitra, P.-C. Musuamba-Kayembe, A. Citenga, J. Tuenakoko-Kulumbula, N. Mapenzi-Kashali, B. Muyembe, J.-C. Makangara-Cigolo, E. Mabika-Bope, K.-M. Kenye, S. Banyima-Satchu, F. Berocan-Underos, B. Bakambu-Nebape, N. Kavugho-Sindani, L. Tibasima-Dhesa, NHLDS consortium, C. Makoha, H. O. Rosette, V. Nakintu, T. Muyigi, G. Pimundu, UVRI consortium, S. Balinandi, D. Ssemwanga, Uganda Ministry of Health consortium, D. Kadobera, D. Euriene, E. Micheal, R. Bahatungire, S. Gidudu, ACE-Uganda consortium, A. Walakira, S. Semawule, K. C. Nabukeera, R. Galiwango, Sentinel consortium, N. Iguosadolo, A. Happi, E. Sijuwola, M. F. Saibu, C. WIlkason, A. Ozonoff, P. Sabeti, H. K. Bosa, D. Mukadi-Bamuleka, C. Olaro, P. Paku-Tshambu, A. Ayouba, S. Mulangu, A. Ssemaganda, N. Loman, A. W. Rimoin, A. Kagiria, S. Wilkinson, B. Lubwama, K. K. Ariën, S. Ahuka-Mundeke, Á. O’Toole, D. Mwamba, C. Happi, M. Peeters, L. Liesenborghs, P. Maes, M. W. Carroll, J. Kindrachuk, P. Akilimali, A. Rambaut, K. Vercauteren, J.-J. Muyembe-Tamfum, T. Wawina-Bokalanga, I. Ssewanyana, P. Mbala-Kingebeni, Emergence of a Bundibugyo virus variant in the 2026 outbreak in the Democratic Republic of the Congo and Uganda. Nat. Med., doi: 10.1038/s41591-026-04628-8 (2026).

4. A. Ayitewala, A. Amuri-Aziza, A. Nsawotebba, T. Wawina-Bokalanga, I. Ssewanyana, S. Nabadda, E. Kinganda-Lusamaki, S. Kanyerezi, V. N. Zalwango, D. Mukadi-Bamuleka, N. Mulopo-Mukanya, G. Luakanda-Ndelemo, A. Kumar, H. K. Bosa, W. Sabiiti, W. Muttamba, B. Bakamutumaho, B. Kirenga, M. Dutt, G. S. Martinez, D. J. Kelvin, C. Olaro, J.-R. O. Aceng, D. Atwiine, P. Mbala-Kingebeni, M. Wayengera, Bundibugyo ebolavirus from the 2026 Ebola outbreak in Uganda and DR Congo: a new variant. The Lancet 408, 317–319 (2026).

5 K. Kupferschmidt, EXCLUSIVE: Congo’s Ebola epidemic started at least 4 months before it was detected (2026). https://www.science.org/content/article/exclusive-congo-s-ebola-epidemic-started-least-4-months-it-was-detected.

6. WHO, Ebola disease caused by Bundibugyo virus - Democratic Republic of the Congo, Disease Outbreak News (2026). https://www.who.int/emergencies/disease-outbreak-news/item/2026-DON614.

7. W. Cao, S. He, G. Liu, H. Schulz, K. Emeterio, M. Chan, K. Tierney, K. Azaransky, G. Soule, N. Tailor, A. Salawudeen, R. Nichols, J. Fusco, D. Safronetz, L. Banadyga, The rVSV-EBOV vaccine provides limited cross-protection against Sudan virus in guinea pigs. Npj Vaccines 8, 91 (2023).

8. D. Falzarano, F. Feldmann, A. Grolla, A. Leung, H. Ebihara, J. E. Strong, A. Marzi, A. Takada, S. Jones, J. Gren, J. Geisbert, S. M. Jones, T. W. Geisbert, H. Feldmann, Single Immunization With a Monovalent Vesicular Stomatitis Virus–Based Vaccine Protects Nonhuman Primates Against Heterologous Challenge With Bundibugyo ebolavirus. J. Infect. Dis. 204, S1082– S1089 (2011).

9. L. E. Hensley, S. Mulangu, C. Asiedu, J. Johnson, A. N. Honko, D. Stanley, G. Fabozzi, S. T. Nichol, T. G. Ksiazek, P. E. Rollin, V. Wahl-Jensen, M. Bailey, P. B. Jahrling, M. Roederer, R. A. Koup, N. J. Sullivan, Demonstration of Cross-Protective Vaccine Immunity against an Emerging Pathogenic Ebolavirus Species. PLoS Pathog. 6, e1000904 (2010).

10. A. Marzi, H. Ebihara, J. Callison, A. Groseth, K. J. Williams, T. W. Geisbert, H. Feldmann, Vesicular Stomatitis Virus–Based Ebola Vaccines With Improved Cross-Protective Efficacy. J. Infect. Dis. 204, S1066–S1074 (2011).

11. C. E. Mire, J. B. Geisbert, A. Marzi, K. N. Agans, H. Feldmann, T. W. Geisbert, Vesicular Stomatitis Virus-Based Vaccines Protect Nonhuman Primates against Bundibugyo ebolavirus. PLoS Negl. Trop. Dis. 7, e2600 (2013).

12. Z. Schiffman, G. Liu, W. Cao, W. Zhu, K. Emeterio, X. Qiu, L. Banadyga, The Ferret as a Model for Filovirus Pathogenesis and Countermeasure Evaluation. ILAR J. 61, 62–71 (2020).

13. R. W. Cross, C. E. Mire, V. Borisevich, J. B. Geisbert, K. A. Fenton, T. W. Geisbert, The Domestic Ferret (Mustela putorius furo) as a Lethal Infection Model for 3 Species of Ebolavirus. J. Infect. Dis. 214, 565–569 (2016).

14. R. Kozak, S. He, A. Kroeker, M.-A. De La Vega, J. Audet, G. Wong, C. Urfano, K. Antonation, C. Embury-Hyatt, G. P. Kobinger, X. Qiu, Ferrets Infected with Bundibugyo Virus or Ebola Virus Recapitulate Important Aspects of Human Filovirus Disease. J. Virol. 90, 9209–9223 (2016).

15. L. Banadyga, G. Wong, X. Qiu, Small Animal Models for Evaluating Filovirus Countermeasures. ACS Infect. Dis. 4, 673–685 (2018).

16. J. Wight, H. Schulz, L. Banadyga, Antibodies Cross-Reactive with Bundibugyo Virus in Ferrets Vaccinated with Ebola Virus Vaccine. Emerg. Infect. Dis. 32, 1360–1363 (2026).

17. R. Burk, L. Bollinger, J. C. Johnson, J. Wada, S. R. Radoshitzky, G. Palacios, S. Bavari, P. B. Jahrling, J. H. Kuhn, Neglected filoviruses. FEMS Microbiol. Rev. 40, 494–519 (2016).

18. K. Kuppalli, R. Lokudu, A. S. Azman, A. Sprecher, G. M. Komanda, I. Ciglenecki, M. Albela, D. Mukendi, Y. Boum, J. B. Nachega, P. Mbala, J.-J. Muyembe-Tamfum, Reconsidering the rVSVΔG-ZEBOV-GP vaccine during the 2026 Bundibugyo virus outbreak. Lancet Infect. Dis., S1473309926003828 (2026).

19. J. R. Andrews, P. K. Mbala, P. K. Mukadi, J. Kindrachuk, N. A. Hoff, A. W. Rimoin, I. I. Bogoch, Ring and community vaccination for Bundibugyo ebolavirus outbreak response: a stochastic network modelling study. medRxiv [Preprint] (2026). 10.64898/2026.07.09.26357654.

20. S. A. Ehrhardt, M. Zehner, V. Krähling, H. Cohen-Dvashi, C. Kreer, N. Elad, H. Gruell, M. S. Ercanoglu, P. Schommers, L. Gieselmann, R. Eggeling, C. Dahlke, T. Wolf, N. Pfeifer, M. M. Addo, R. Diskin, S. Becker, F. Klein, Polyclonal and convergent antibody response to Ebola virus vaccine rVSV-ZEBOV. Nat. Med. 25, 1589–1600 (2019).

21. M. Halbrook, S. Merritt, N. A. Hoff, P. Mukadi, J. P. Kompany, K. Musene, M. Beya, H. Kalengi, M. Tambu, J. D. Kelly, A. H. Ball, A. To, C. W. Woods, B. P. Nicholson, M. T. McClain, T. Wong, L. E. Hensley, J. Kindrachuk, A. T. Lehrer, P. Mbala, A. W. Rimoin, Longitudinal Bundibugyo Virus Glycoprotein Seroreactivity Following rVSVΔG-ZEBOV-GP Vaccination in Outbreak-Affected Populations of the Democratic Republic of the Congo. medRxiv [Preprint] (2026). 10.64898/2026.06.22.26356273.

22. E. Lhomme, A. Wiedemann, A. Ayouba, S. Ben-Farhat, G. Thaurignac, C. Roy, A. H. Beavogui, S. Doumbia, M. Kieh, B. Leigh, S. Sow, S. A. Migueles, D. Watson-Jones, Y. Yazdanpanah, R. Thiébaut, M. Peeters, L. Richert, Y. Levy, PREVAC Study Team, Cross-Reactive Bundibugyo Antibody Responses after Receipt of Licensed Ebola Vaccines. N. Engl. J. Med., doi: 10.1056/NEJMc2608018 (2026).

23. P. LaRochelle, M. McKnight, U. U. Patrick, M. A. Davin, Congolese hospital staff cohort admitted with infectious symptoms in the setting of the Bundibugyo virus outbreak, April and May 2026, Bunia, DR Congo. Lancet Infect. Dis., S1473309926004202 (2026).

24. R. F. Grais, S. B. Kennedy, B. E. Mahon, S. A. Dubey, R. J. Grant-Klein, K. Liu, J. Hartzel, B.-A. Coller, C. Welebob, M. E. Hanson, J. K. Simon, Estimation of the correlates of protection of the rVSVΔG-ZEBOV-GP Zaire ebolavirus vaccine: a post-hoc analysis of data from phase 2/3 clinical trials. Lancet Microbe 2, e70–e78 (2021).

25. S. M. Jones, H. Feldmann, U. Ströher, J. B. Geisbert, L. Fernando, A. Grolla, H.-D. Klenk, N. J. Sullivan, V. E. Volchkov, E. A. Fritz, K. M. Daddario, L. E. Hensley, P. B. Jahrling, T. W. Geisbert, Live attenuated recombinant vaccine protects nonhuman primates against Ebola and Marburg viruses. Nat. Med. 11, 786–790 (2005).

26. A. Marzi, F. Engelmann, F. Feldmann, K. Haberthur, W. L. Shupert, D. Brining, D. P. Scott, T. W. Geisbert, Y. Kawaoka, M. G. Katze, H. Feldmann, I. Messaoudi, Antibodies are necessary for rVSV/ZEBOV-GP–mediated protection against lethal Ebola virus challenge in nonhuman primates. Proc. Natl. Acad. Sci. 110, 1893–1898 (2013).

27. A. C. Shurtleff, J. C. Trefry, S. Dubey, M. M. E. Sunay, K. Liu, Z. Chen, M. Eichberg, P. M. Silvera, S. A. Kwilas, J. W. Hooper, S. Martin, J. K. Simon, B.-A. G. Coller, T. P. Monath, rVSVΔG-ZEBOV-GP Vaccine Is Highly Immunogenic and Efficacious Across a Wide Dose Range in a Nonhuman Primate EBOV Challenge Model. Viruses 17, 341 (2025).

28. G. Wong, J. S. Richardson, S. Pillet, A. Patel, X. Qiu, J. Alimonti, J. Hogan, Y. Zhang, A. Takada, H. Feldmann, G. P. Kobinger, Immune Parameters Correlate with Protection Against Ebola Virus Infection in Rodents and Nonhuman Primates. Sci. Transl. Med. 4 (2012).

29. N. A. Kuzmina, P. Younan, P. Gilchuk, R. I. Santos, A. I. Flyak, P. A. Ilinykh, K. Huang, N. M. Lubaki, P. Ramanathan, J. E. Crowe, A. Bukreyev, Antibody-Dependent Enhancement of Ebola Virus Infection by Human Antibodies Isolated from Survivors. Cell Rep. 24, 1802–1815.e5 (2018).

30. P. Fletcher, K. L. O’Donnell, F. Feldmann, J. F. Rhoderick, C. S. Clancy, J. A. Haase, C. A. Prator, B. J. Smith, B. M. Gunn, H. Feldmann, A. Marzi, Fast-acting single-dose vesicular stomatitis virus-Sudan virus vaccine: a challenge study in macaques. Lancet Microbe 6, 101244 (2025).

31. WHO, “Third meeting of the WHO Technical Advisory Group on candidate vaccine prioritization (TAG-CVP) for Bundibugyo virus disease outbreak response: meeting report” (Geneva, 2026); 10.2471/B09865.

32. A. M. Henao-Restrepo, A. Camacho, I. M. Longini, C. H. Watson, W. J. Edmunds, M. Egger, M. W. Carroll, N. E. Dean, I. Diatta, M. Doumbia, B. Draguez, S. Duraffour, G. Enwere, R. Grais, S. Gunther, P.-S. Gsell, S. Hossmann, S. V. Watle, M. K. Kondé, S. Kéïta, S. Kone, E. Kuisma, M. M. Levine, S. Mandal, T. Mauget, G. Norheim, X. Riveros, A. Soumah, S. Trelle, A. S. Vicari, J.-A. Røttingen, M.-P. Kieny, Efficacy and effectiveness of an rVSV-vectored vaccine in preventing Ebola virus disease: final results from the Guinea ring vaccination, open-label, cluster-randomised trial (Ebola Ça Suffit!). The Lancet 389, 505–518 (2017).

33. M. Garbutt, R. Liebscher, V. Wahl-Jensen, S. Jones, P. Möller, R. Wagner, V. Volchkov, H.-D. Klenk, H. Feldmann, U. Ströher, Properties of Replication-Competent Vesicular Stomatitis Virus Vectors Expressing Glycoproteins of Filoviruses and Arenaviruses. J. Virol. 78, 5458– 5465 (2004).

34. Qiagen, ssRNA Mass Moles Converter (2026). https://www.qiagen.com/us/applications/enzymes/tools-and-calculators/rna-mass-to-moles-conventer.

35. L. J. Reed, H. Muench, A simple method of estimating fifty per cent endpoints. Am. J. Hyg. 27, 493–497 (1934).

36. R Core Team, R: A language and environment for statistical computing., R Foundation for Statistical Computing (2025); https://www.R-project.org/.

37. T. M. Therneau, A Package for Survival Analysis in R, version 3.8-11 (2026); https://CRAN.R-project.org/package=survival.

38. C. Wilke, B. Wiernik, ggtext: Improved Text Rendering Support for ’ggplot2, version 0.1.2 (2022); https://CRAN.R-project.org/package=ggtext.

39. T. L. Pedersen, patchwork: The Composer of Plots, version 1.3.2 (2025); https://CRAN.R-project.org/package=patchwork.

40. A. Kassambara, rstatix: Pipe-Friendly Framework for Basic Statistical Tests, version 0.7.3 (2025); https://CRAN.R-project.org/package=rstatix.

41. H. Wickham, T. L. Pedersen, D. Seidel, scales: Scale Functions for Visualization, version 1.4.0 (2025); https://CRAN.R-project.org/package=scales.

42. H. Wickham, M. Averick, J. Bryan, W. Chang, L. McGowan, R. François, G. Grolemund, A. Hayes, L. Henry, J. Hester, M. Kuhn, T. Pedersen, E. Miller, S. Bache, K. Müller, J. Ooms, D. Robinson, D. Seidel, V. Spinu, K. Takahashi, D. Vaughan, C. Wilke, K. Woo, H. Yutani, Welcome to the Tidyverse. J. Open Source Softw. 4, 1686 (2019).

