## Supplementary Material for "rVSV-EBOV vaccination protects ferrets from lethal Bundibugyo virus disease"

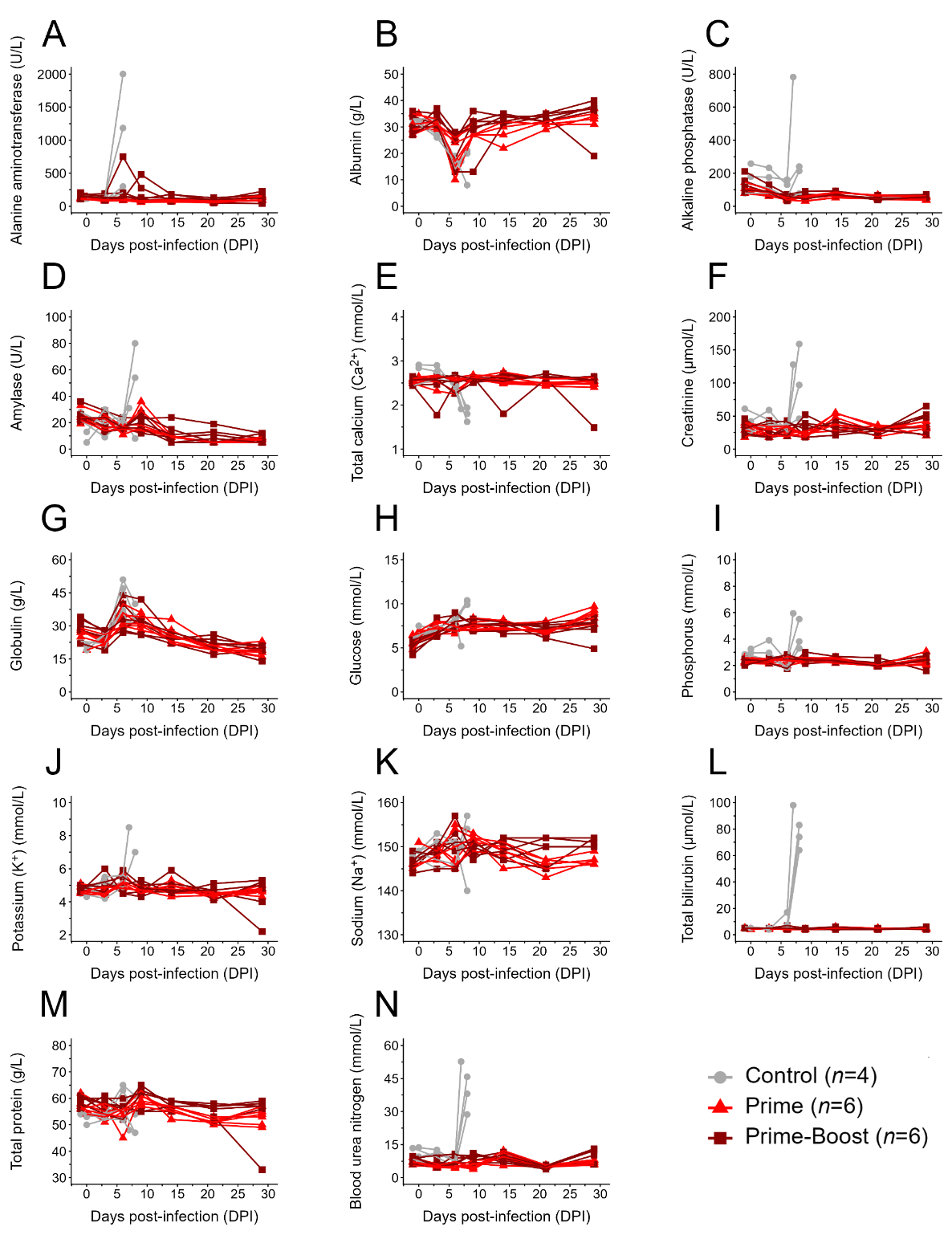

**Fig. S1.** | **Blood biochemistry parameters in all ferrets.** Alanine aminotransferase (**A**), albumin (**B**), alkaline phosphatase (**C**), amylase (**D**), total calcium (**E**), creatinine (**F**), globulin (**G**), glucose (**H**), phosphorus (**I**), potassium (**J**), sodium (**K**), total bilirubin (**L**), total protein (**M**), and blood urea nitrogen (**N**) levels were assessed in the blood of all ferrets after BDBV inoculation.

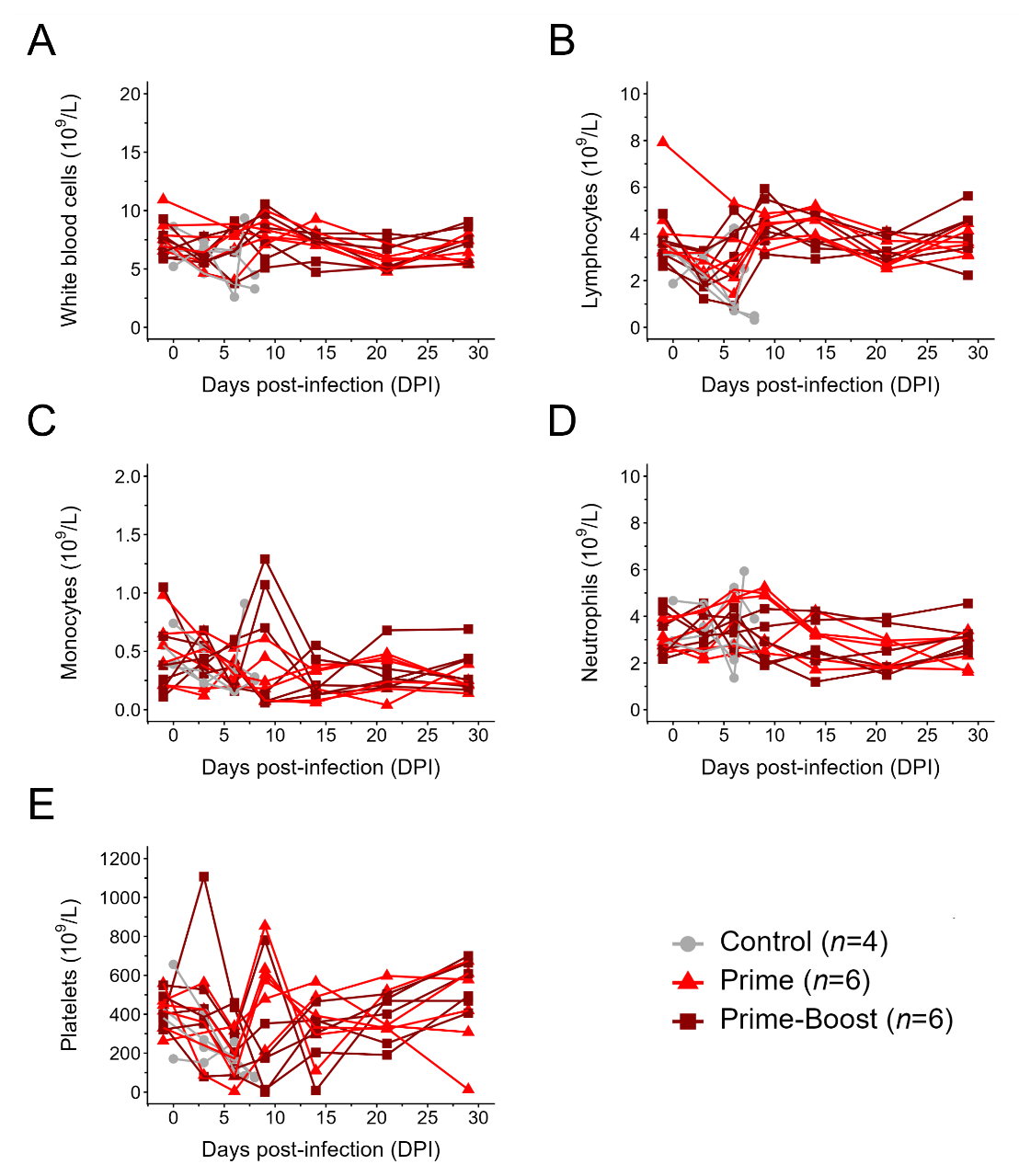

**Fig. S2.** | **Complete blood cell counts in all ferrets.** White blood cells (**A**), lymphocytes (**B**), monocytes (**C**), neutrophils (**D**), and platelets (**E**) were enumerated in the blood of all ferrets after BDBV inoculation.

| **Table S1. Animal Information and Disease Outcome** | | | | | |
| --- | --- | --- | --- | --- | --- |
| **Group** | **Vaccine** | **ID** | **Sex** | **Disease Manifestation** | **Outcome** |
| 1 | rVSV-EBOV Prime | 642 | M | Elevated temperature on 5 DPI only; otherwise normal | Survived |
|  |  | 570 | M | No clinical signs observed | Survived |
|  |  | 502 | M | No clinical signs observed | Survived |
|  |  | 158 | F | Elevated temperature on 6 DPI only; otherwise normal | Survived |
|  |  | 140 | F | No clinical signs observed | Survived |
|  |  | 069 | F | Elevated temperature on 3 DPI only; otherwise normal | Survived |
| 2 | rVSV-EBOV Prime-Boost | 499 | M | No clinical signs observed | Survived |
|  |  | 405 | M | Elevated temperature on 5, 6 DPI with increased respiration on 6 DPI | Survived |
|  |  | 324 | M | No clinical signs observed | Survived |
|  |  | 051 | F | Elevated temperature on 6 DPI only; otherwise normal | Survived |
|  |  | 000 | F | No clinical signs observed | Survived |
|  |  | 674 | F | No clinical signs observed | Survived |
| 3 | Control/  Saline | 126 | M | Elevated temperature beginning on 6 PDI; severe disease culminating in euthanasia on 8 DPI; petechial rash | Died |
|  |  | 255 | F | Elevated temperature beginning on 6 DPI; severe disease culminating in euthanasia on 8 DPI | Died |
| 4 | Staggered Control | 477 | M | Elevated temperature beginning on 5 DPI; severe disease culminating in euthanasia on 7 DPI; petechial rash and evidence of hemorrhage | Died |
|  |  | 322 | F | Elevated temperature beginning on 5 DPI; severe disease culminating in euthanasia on 8 DPI; evidence of hemorrhage | Died |
